# Individual-based simulations of genome evolution with ancestry: the GenomeAdmixR R package

**DOI:** 10.1101/2020.10.19.343491

**Authors:** Thijs Janzen, Fernando Diaz

**Affiliations:** Groningen Institute for Evolutionary Life Sciences, University of Groningen, Box 111039700 CC Groningen, The Netherlands; Carl von Ossietzky University, Carl-von-Ossietzky-Str. 9–11, 26111, Oldenburg, Germany; Department of Entomology, University of Arizona, Tucson, AZ, United States of America

**Keywords:** Genome evolution, admixture, ancestry, individual-based modeling, Evolve and Resequence

## Abstract

1. Hybridization between populations or species results in a mosaic of the two parental genomes. This and other types of genome admixture have received increasing attention for their implications in speciation, human evolution, Evolve and Resequence (E&R) and genetic mapping. However, a thorough understanding of how local ancestry changes after admixture, and how selection affects patterns of local ancestry remains elusive. The complexity of these questions limits analytical treatment, but these scenarios are specifically suitable for simulation.
2. Here, we present the R package GenomeAdmixR, which uses an individual-based model to simulate genomic patterns following admixture forward in time. GenomeAdmixR provides user-friendly functions to set up and analyze simulations under evolutionary scenarios with selection, linkage and migration.
3. We show the flexible functionality of the GenomeAdmixR workflow by demonstrating 1) how to design an E&R simulation using GenomeAdmixR and 2) how to use GenomeAdmixR to verify analytical expectations following from the theory of junctions.
4. GenomeAdmixR provides a mechanistic approach to explore expected genome responses to realistic admixture scenarios. With this package, we aim to aid researchers in testing specific hypotheses based on empirical findings involving admixing populations.

## INTRODUCTION

Genetic exchange has long been recognized as an important driver of genetic diversity, from the recombination of conspecific genomes and the evolution of speciation with migration (Abbott *et al*. 2013) to introgressive hybridization (Janzen *et al*. 2018; Lavretsky *et al*. 2019) and horizontal gene transfer (Keeling & Palmer 2008). With recent advances in sequencing technologies, the view of genetic exchange has now moved from information at specific loci (*e*.*g*. alleles, genes) to the level of whole genome admixture (Leitwein *et al*. 2018; Lavretsky *et al*. 2019; Chafin & Douglas 2020). The focus is now on understanding how major drivers of evolution (*i*.*e*. recombination, selection and migration) shape the linear connection of loci along the genome (*i*.*e*. synteny and linkage) and how these processes evolve under particular scenarios of genome divergence. Understanding these dynamics has helped to elucidate the impact of migration on human evolution (Hellenthal *et al*. 2014; Payseur & Rieseberg 2016), the impact of hybridization on speciation (Schumer *et al*. 2014b, 2018), and helped as well in designing experiments relying on admixture, for instance Evolve and Resequence (E&R) experiments (Franssen *et al*. 2017; Barghi & Schlötterer 2019; Otte & Schlötterer 2020).

Recombination is one of the main drivers shaping macro-genomic patterns after admixture, where contiguous blocks of ancestry are broken down into smaller blocks over time (Fisher 1954, 1959). As time progresses, genomes transform into macro-genomic mosaics consisting of many small contiguous ancestry blocks. Our understanding of how these mosaics form is relatively robust, but analytical treatments are only known for boundary cases, such as admixture between two distant populations, or for restricted backcrossing (Fisher 1954, 1959; Stam 1980; Macleod *et al*. 2005; Janzen *et al*. 2018; Lavretsky *et al*. 2019).

Linkage Disequilibrium blocks can also be generated by selection through genetic hitchhiking or selection of epistatic interactions, which can alter genome-wide patterns of genetic variation, population dynamics and evolvability (Gerrish *et al*. 2007; Arnold & Kunte 2017; Zhou *et al*. 2017). Thus, macro-genomic patterns can be driven both by recombination and selection alike. However, a mathematical treatment of the interaction between selection, recombination and admixture is currently lacking. In contrast, individual-based simulations are very suitable to explore these interactions, for instance using packages such as `MimicrEE2` (Vlachos & Kofler 2018) (written in Java) and `SLiM` (Haller & Messer 2017, 2018) (written in C++). Unfortunately, these do not explicitly track local ancestry, and information on haplotypes is often only indirectly available or requires extensive post-processing. `forqs` (Kessner & Novembre 2014) and `SELAM` (Corbett-Detig & Jones 2016) (both written in C++) remedy this issue and are specifically focused on tracking local ancestry and inheritance of ancestry blocks, however these lack ease of use (requiring for instance local compilation of C++ code, and configuration of input files) and are not easily applied cross-platform. The R package `plmgg` (Cottin *et al*. 2020) is easily used cross-platform, but focuses mainly on plant-like admixture, including selfing.

Here, we present GenomeAdmixR, a software package written in the R language. The R language is readily used within biology and can easily be used cross-platform. GenomeAdmixR provides routines to simulate admixture in a host of scenarios (Figure 1), and includes a wide range of routines available to analyze and visualize simulation results.

**Figure 1.** Overview of applications of the GenomeAdmixR package. The GenomeAdmixR package can be used to **A**) simulate admixture of genomes (rectangular, colored) over time, passed on between individuals (rounded squares) when two populations (dotted lines) meet. **B**) Simulate ongoing genetic exchange under migration, **C**) explore the impact of selection for alleles from the red ancestor in the first section of the genome and **D**) Simulate the formation of Recombinant Inbred Lines, where continued inbreeding generates fully homozygous.

## MATERIALS and METHODS

### Description

The interface of GenomeAdmixR is written in the R programming language (Team 2020), with the underlying simulation code using C++, integrated using Rcpp (Eddelbuettel & Francois 2011). Similar to SliM (Haller & Messer 2017, 2018) and SELAM (Corbett-Detig & Jones 2016), GenomeAdmixR simulates a population forward in time usidng a Wright-Fisher model with non-overlapping generations and a constant population size. GenomeAdmixR simulates diploid individuals that sexually reproduce. For computational tractability, only one pair of chromosomes is simulated, and all individuals are assumed to be hermaphroditic. Recombination is modeled in the same way as in for instance SLiM and SELAM (Corbett-Detig & Jones 2016; Haller & Messer 2018), *i*.*e*. as a Poisson process, with the number of crossovers Poisson distributed with the size of the chromosome in Morgan as rate parameter. The location of crossovers is drawn from a uniform distribution, without interference.

The package offers two different implementations of this simulation scheme. Firstly, the user can use the ***Ancestry module*** of GenomeAdmixR to perform the simulations using known local ancestry, where the simulation tracks the locations of changes in local ancestry (like SELAM and forqs) but does not explicitly model nucleotides or mutation (as mutation between ancestries complicates the simplifying assumptions of junctions’ theory used to propagate the simulation). Secondly, the *Sequence module* can be used to perform simulations starting with sequencing data. Recombination is provided in cM/Mb and simulations can therefore be performed on a section rather than the entire chromosome. When using the ***Sequence module***, mutation options are included as well. Sequencing data can be loaded from VCF or PLINK format (using the function `read_input_data`). To demonstrate the functionality of the package, we have included sample data from the *Drosophila melanogaster* Reference Panel (MacKay *et al*. 2012; Huang *et al*. 2014). The sample data consists of 5000 SNPs with minimal allele frequency of 0.05, located along the 3R chromosome arm.

### Usage

#### The GenomeAdmixR package can be installed from CRAN

`> install.packages`(“`GenomeAdmixR`”)

The core of GenomeAdmixR is formed by the function `simulate_admixture`, customizable with two modules: the ***Ancestry module*** and the ***Sequence module*** (Table 1). Here we show how to implement simulations using both modules side-by-side. To simulate a simple admixture scenario where two distinct populations hybridize (e.g. a population of admixed individuals resulting from a single mating event between two unrelated individuals, resulting in an exactly 50/50 mixing of ancestral genomes in the first generation) to form a population, which continues to admix for 100 generations, we write:

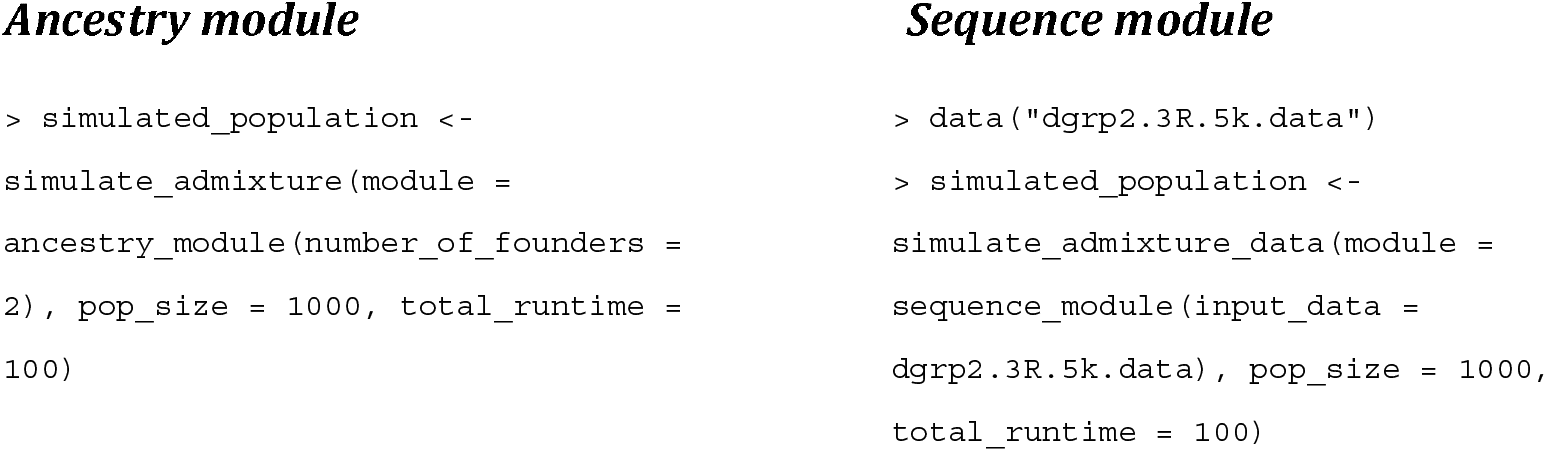

**Table 1.**
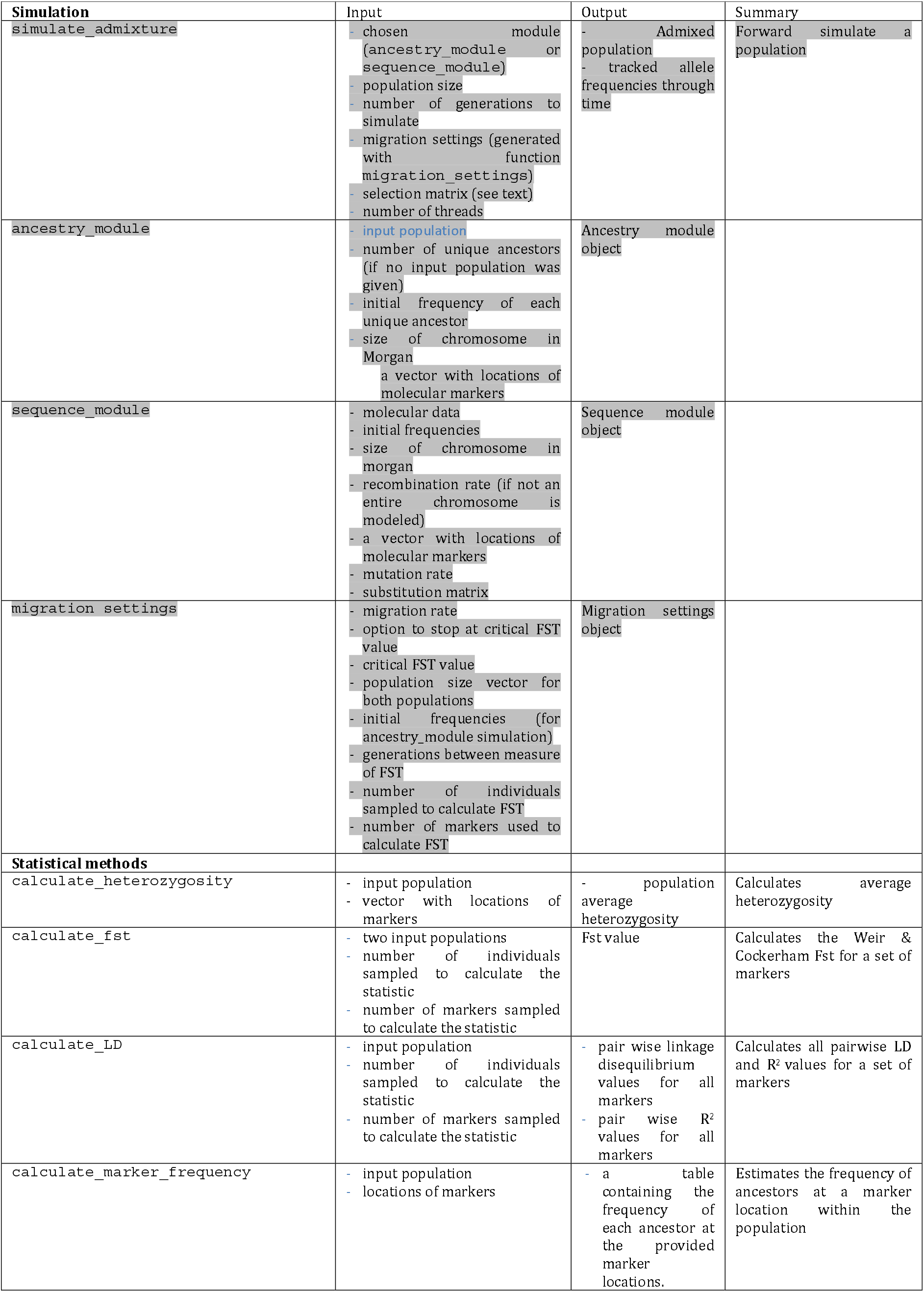
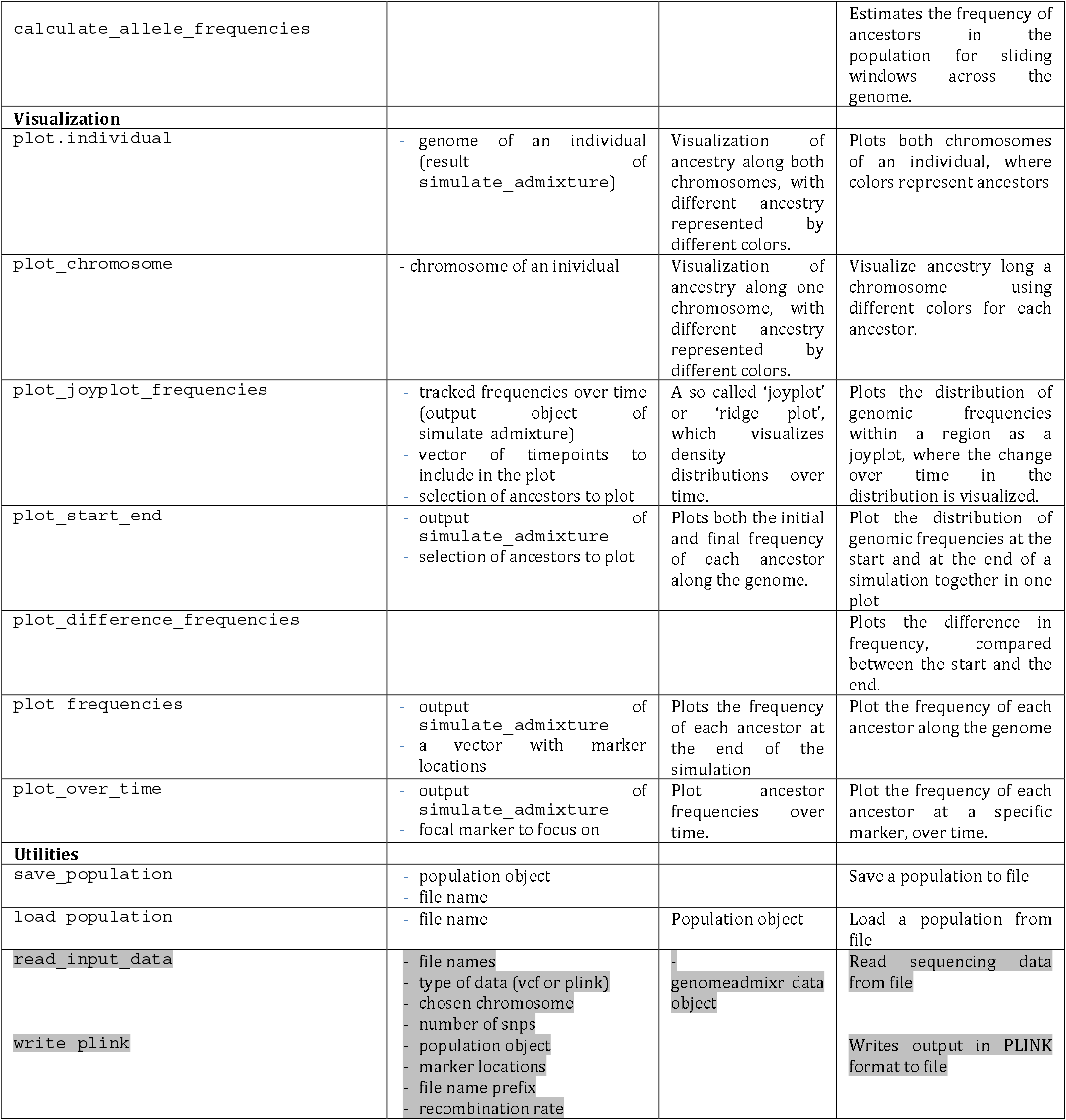
Overview of available functions in the package

#### Molecular markers and statistic estimations

GenomeAdmixR includes the functionality to track molecular markers located along the genome, in line with molecular methods (Schumer *et al*. 2014a; Dennenmoser *et al*. 2019; Lavretsky *et al*. 2019). For the *ancestry module*, markers also serve to simulate the uncertainty in tracking local ancestry due to limited coverage. The markers are assumed to be on a fixed position (in Morgan for the *ancestry module*, and in bp for the *sequence module*) and are tracked over time. Resulting local information is returned from the function in long format, facilitating downstream analyses. Repeating the previous scenario, but now tracking 1000 markers at fixed positions along the genome:

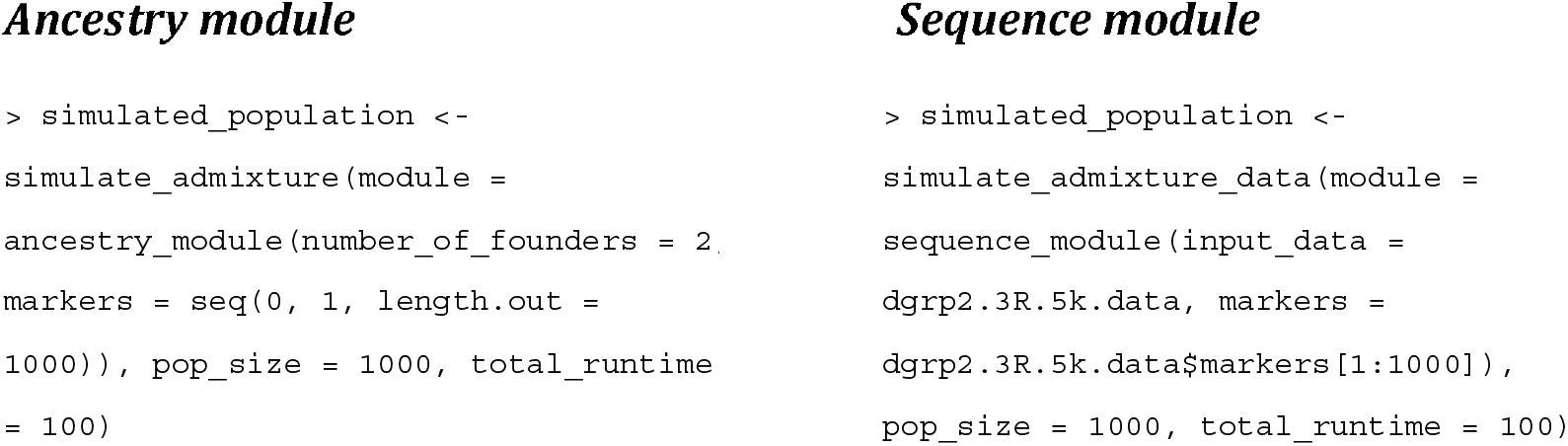

Based on these markers, a set of genetic diversity statistics can be calculated, including allele frequency (`calculate_allele_frequencies`, Table 1), heterozygosity (`calculate_heterozygosity,` Table 1), linkage disequilibrium (`calculate_ld`, Table 1), and F_ST_ (F_ST_ is estimated following Weir and Cockerham (Weir & Cockerham 1984), using the R package `hierfstat` (Goudet 2005) for the implementation, see `calculate_fst`, Table 1), as well as changes of these over generations for better assessing significance of simulated evolution. When using the *ancestry module*, caution should be taken when using more than four distinct ancestries, because most genetic diversity statistics assume a maximum nucleotide number of four.

#### Demonstration of genetic diversity statistics

We estimate LD and average heterozygosity at different points in time. Our expectation is for LD and heterozygosity to decrease over time. In the code below, we either use 100 regularly spaced markers in Morgan (*ancestry module*), or we draw 100 random marker positions from the data (*sequence module*).

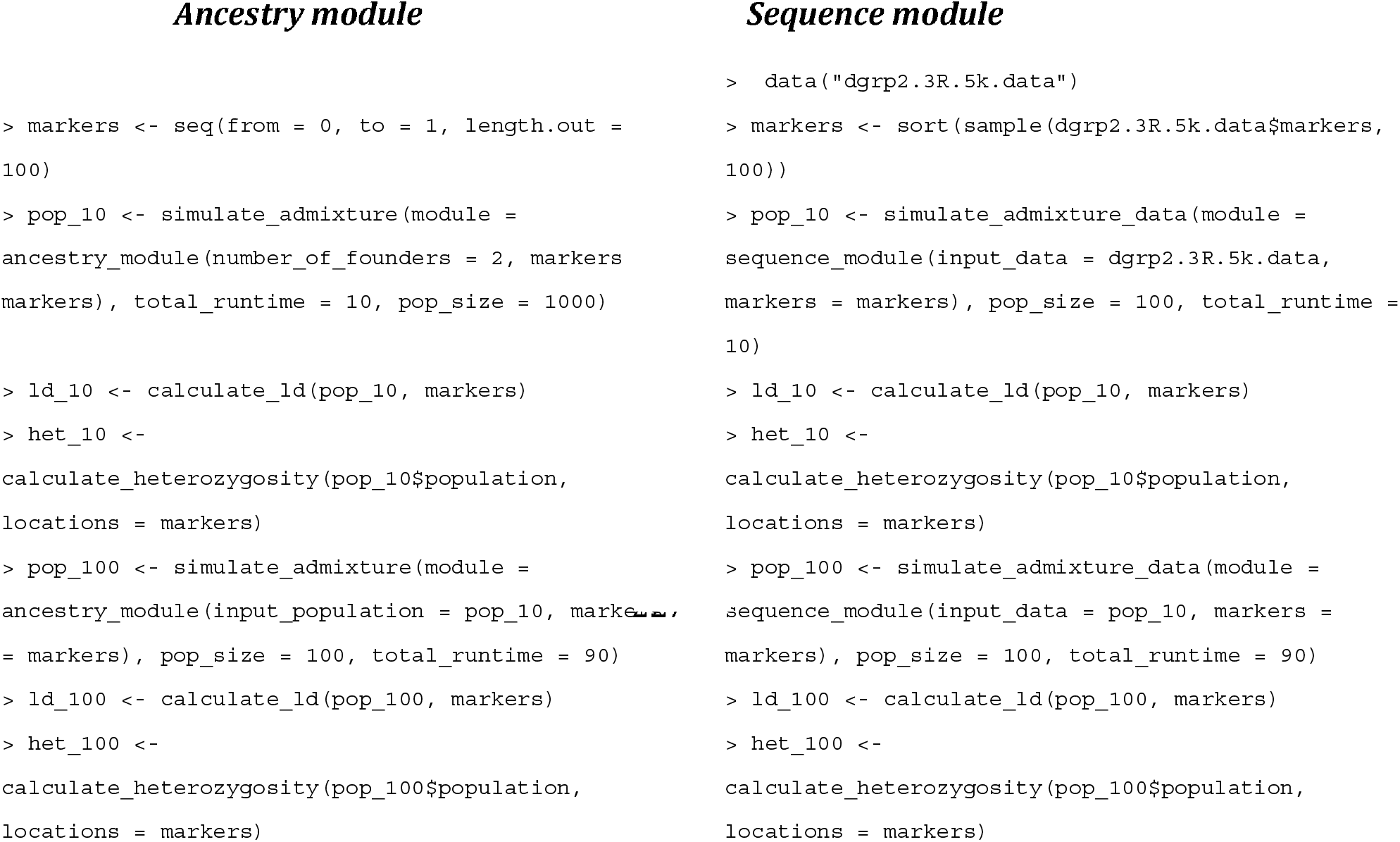

The results (Figure 2) show that LD (here plotted as the correlation (R^2^) between loci) initially is relatively high, especially for closely located loci. However, as time increases, LD disappears. Similarly, initially heterozygosity is high (as expected), but decreases over time across the genome, until most markers are fixed. Patterns are similar across the *ancestry* and *sequence* simulations, although initially, LD is slightly higher for the *sequence* simulations..

**Figure 2.**
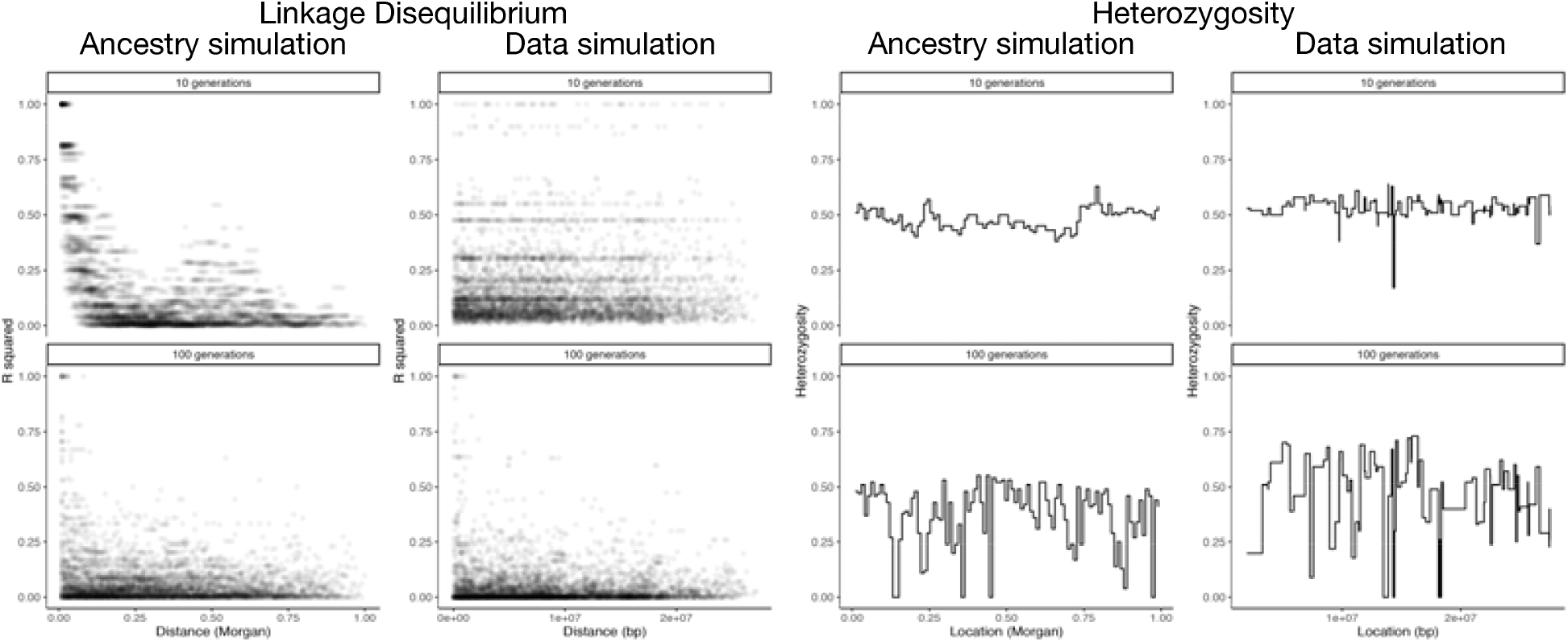
Plot showing Linkage Disequilibrium (R^2^) (left) and average heterozygosity (right) for a population 10 and 100 generations after admixture, where the initial admixture event involved two unrelated populations. Shown are results using the *ancestry module* (left columns) and the *sequence module* (right columns).

#### Visualization

`simulate_admixture` by default returns the frequencies of all ancestors at the start and the end of the simulation, if markers are specified. Allele frequency changes can be visualized using `plot_difference_frequencies_` (Table 1, Figure 3) or `plot_start_end` (Table 1, Figure 3) and using `plot_frequencies` (Table 1, Figure 3) the final frequencies are plotted. The functions `plot_over_time` (Table 1, Figure 3) and `plot_joyplot_frequencies` (Table 1, Figure 3) provide functionality to show allele frequency changes over time. All plotting functions return ggplot2 (Wickham 2009) objects, which can be further customized by the user.

**Figure 3.**
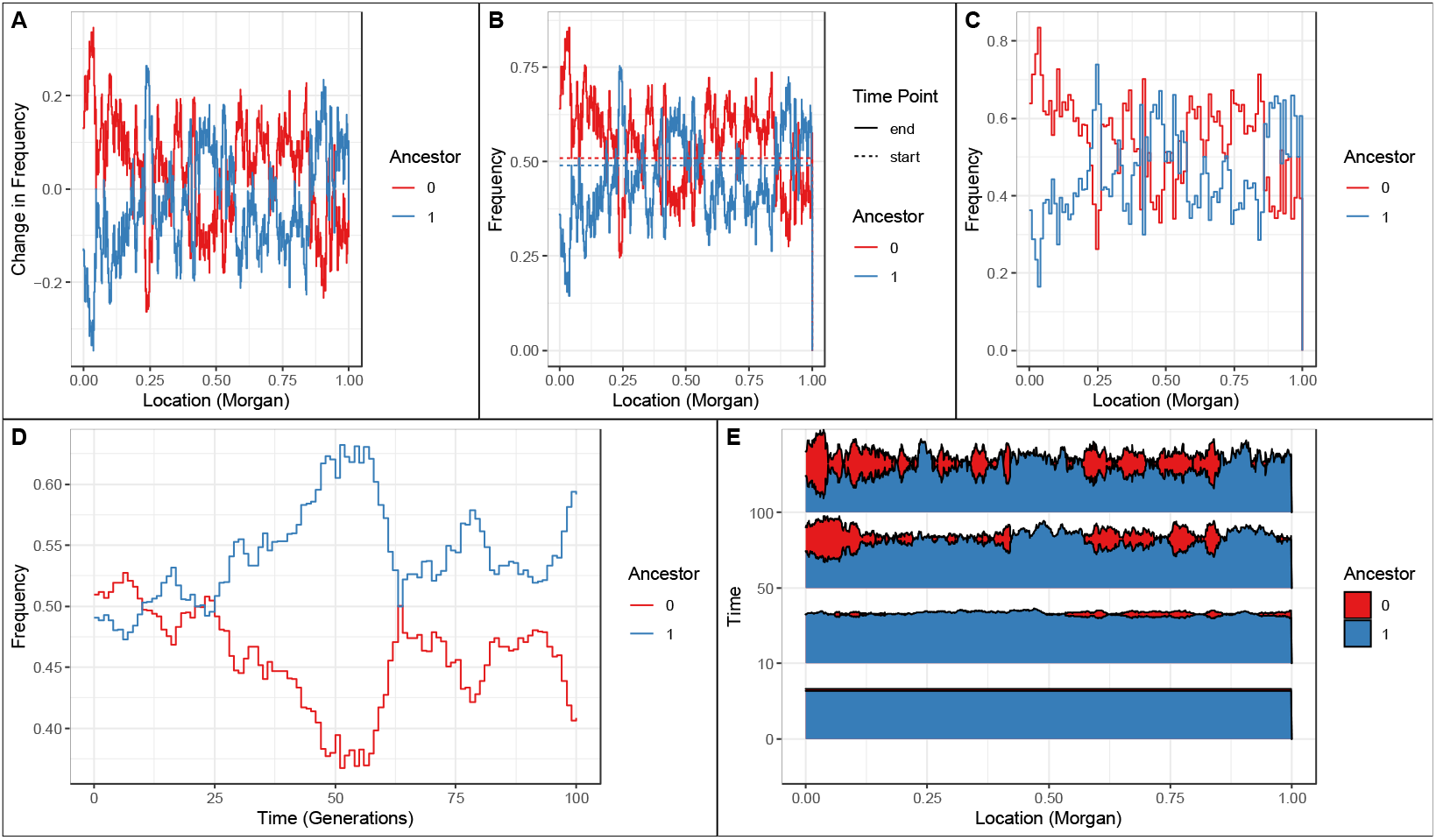
Example visualizations available in GenomeAdmixR. Shown are example plots after an hybridization scenario using the *ancestry module* between two distinct ancestral populations, parameter values used are: population size = 10,000, total runtime = 100. 1000 markers evenly spaced in [0, 1] were used. **A**: `plot_difference_frequencies`, which visualizes the average change in frequency between the start and end of the simulation **B**: plot_start_end, which plots the average frequencies per ancestor, both at the start and the end of the simulation. **C:** plot_frequencies, which plots the average frequency of each ancestor across the genome **D:** plot_over_time, which plots the average frequency of a single marker (here a marker at 50 cM) in the population, over time. **E:** plot_joyplot_frequencies, which visualizes the average frequency over time.

#### Migration

`Simulate_admixture` can be extended further by including migration, where the specific settings for migration can be specified using the function ‘`migration_settings`’ (Table 1). Migration is simulated by evolving two independent populations in parallel, given a fraction migrants each generations.

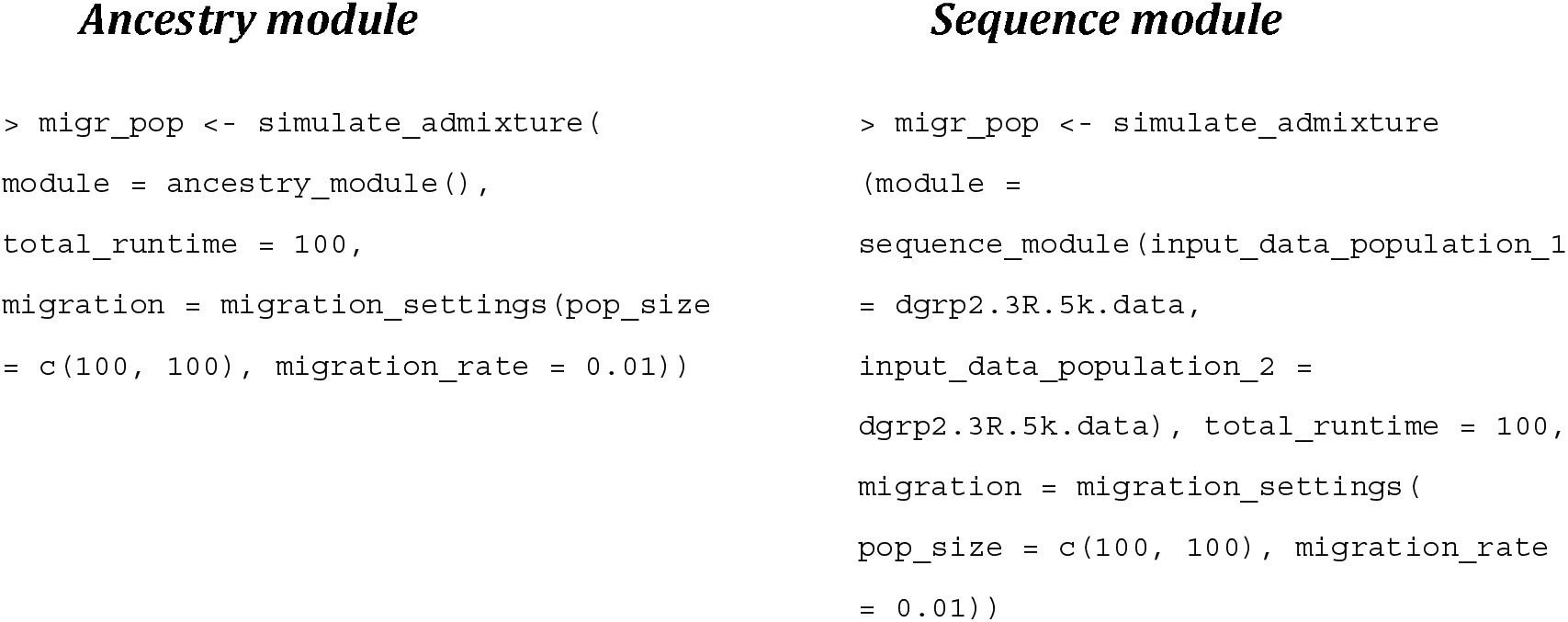

Then, we can analyze genetic divergence across these two populations using the F_ST_ statistic:

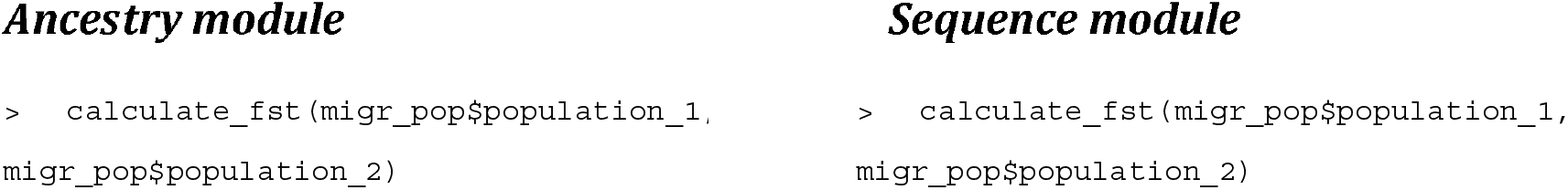

There is functionality included to stop the simulation once a threshold of genetic differentiation (*i*.*e. F*_ST_) between the two populations is reached, which facilitates scenarios where two related, but genetically different populations are required.

#### Selection

`simulate_admixture` provides options to impose fitness benefits upon individuals that contain a marker under selection (where selection is interpreted as favouring alleles from a specific ancestor (*ancestry module*) or SNP (*sequence module*). We follow conventional fitness notation (Crow & Kimura 1970) and denote [w_aa_, w_Aa_, w_AA_] = [*1, 1*+*hs, 1*+*s*], where *w* indicates fitness, *s* indicates the selective benefit of allele *A*, and *h* indicates the degree of dominance. Parents of offspring in the next generation are drawn from the parental population proportional to fitness.

The selection matrix [location, w_aa_, w_Aa_, w_AA_, ancestor] includes information on the selected marker, the different fitness weights, and the origin of the allele under selection. The weights can be freely provided by the user, allowing for the implementation of a wide range of potential selective benefits upon receiving a marker, including overdominance and epistasis.

Furthermore, the user is not restricted to providing selection on a single marker and can expand the selection matrix by adding extra rows for each marker. This opens up the possibility for the user to explore polygenic selection (as for instance for a Quantitative Trait Locus), by specifying several loci with defined distances, where each locus provides a small fitness benefit. Hence, the focal trait under selection here is translated directly in its resulting fitness effect. If multiple loci are under selection, the fitness of an individual is the product of the finesses at each locus. An example of adding selection on two markers at locations of 50 and 60 cM to the previously selected hybridization scenario is:

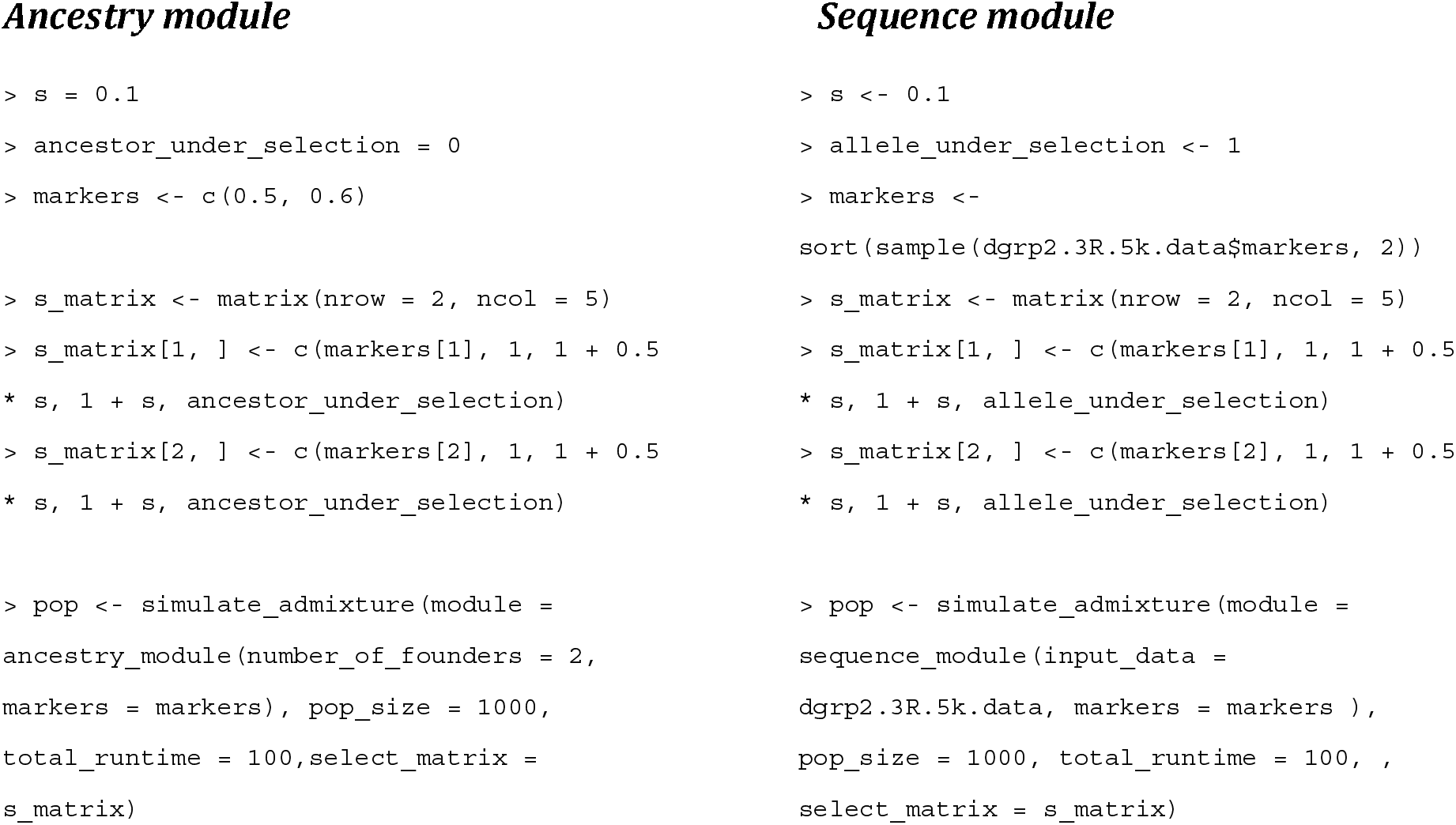

### Demonstration of the package

To demonstrate the functionality of GenomeAdmixR, we firstly show a series of simulations to compare alternative strategies in Evolve & Resequence (*E*&*R*) experiments. Secondly, we show how simulations of GenomeAdmixR match analytical predictions in junction theory.

#### E&R experiment

One of the goals in *E&R* experiments is to identify underlying genetic loci driving responses to selection (Burke & Rose 2009; Schlötterer *et al*. 2015). The most common sampling method starts with a “w*ell mixed population*” founded by individuals sampled from the same natural population (*Model 1*). This method minimizes Linkage Disequilibrium (*LD*) while trying to compensate for genetic variability by performing large sampling (Kofler & Schlötterer 2014). Alternatively, genetic variability can be maximized by sampling from different populations, but this can generate additional *LD* (Kawecki *et al*. 2012; Kofler & Schlötterer 2014; Schlötterer *et al*. 2015). However, it has been proposed that genetic bases of population’ or species’ differences can be investigated by investigating mixing lines from different populations or species (Parts *et al*. 2011). This method (*Model 2*) has been used in yeast to fine-tune Quantitative Trait Locus (*QTL)* studies from phenotypically extreme inbred lines using experimental selection (Parts *et al*. 2011; Koide *et al*. 2012).

We used GenomeAdmixR to simulate the entire *E&R* pipeline and explore the impact of both sampling schemes (Figure S1 demonstrates the simulation setup). Figure 4 (for a single simulation) indicates that the resolution to detect the marker under selection in the experiment was much higher in *Model 1*. Also, *Model 1* seems to experience less genetic hitchhiking compared to *Model 2*. We performed 10 independent simulations with the same results, and replicates can be used for further analysis and statistical tests. Overall, the obtained results indicate that *Model 1* is for this pipeline the preferred approach. We implemented the mentioned pipeline using both the *ancestry module* (Figure 4a) and the *sequence module* (Figure 4b, using *D. melanogaster* Reference Panel data of the 3R arm chromosome) rendering largely congruent results.

**Figure 4.**
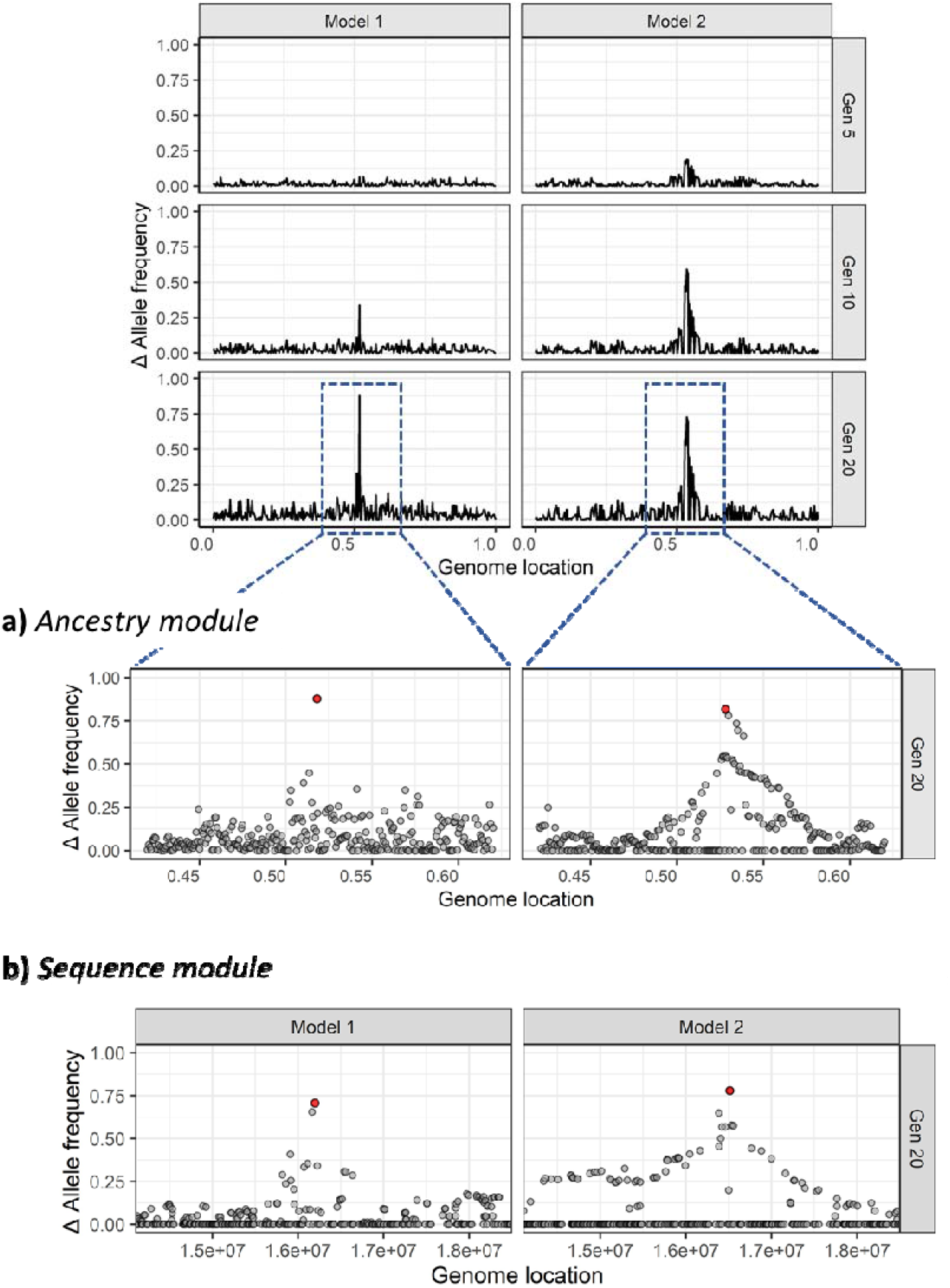
Summary of results obtained after simulating two scenarios for an *E&R* experiment using the GenomeAdmixR package. Genome-wide ⍰⍰allele frequency after 5, 10 and 20 generations of selection are compared between the two models. The models differ in their starting populations as they were founded form a single or diverging populations (*e*.*g. Models 1* vs *Model 2* respectively). Model 1, founded by 100 isofemales form a single population; Model 2, founded by two isofemales form diverging populations. Selection was simulated for only one location (red point in the lower zoomed panel) in a 0.2 Morgan window, and tracked in 1000 markers, while the rest of the genome evolves neutrally. The results are shown for simulations using the **a)** *Ancestry module* and, **b)** the *Sequence module* starting with data from the D. melanogaster Reference Panel.

We further expanded *Model 1*, using individuals instead of isofemale lines (*Models 3* and 4) revealing similar trends (See Figure S2).

#### Theory of junctions

From the extended theory of junctions (Janzen *et al*. 2018) we know that the expected number of junctions (where a junction delineates the end of one contiguous stretch of genomic content from the same ancestor, and the start of another), depends on the initial heterozygosity H_0_ (where heterozygosity here reflects a locus with two alleles stemming from two different ancestors, e.g. hetero-ancestry). Janzen’s extended theory of junctions focuses specifically on the scenario with two ancestors, whereas GenomeAdmixR allows for an arbitrary number of ancestors. We simulate the accumulation of junctions using the *ancestry module* of GenomeAdmixR for two population sizes (100 and 1000 individuals) and vary the number of ancestors *n* in [2, 4, 8]. We find that when we average the number of accumulated junctions over 1000 replicates, the average number of junctions closely follows that of our analytical expectation (Figure 5).

**Figure 5.**
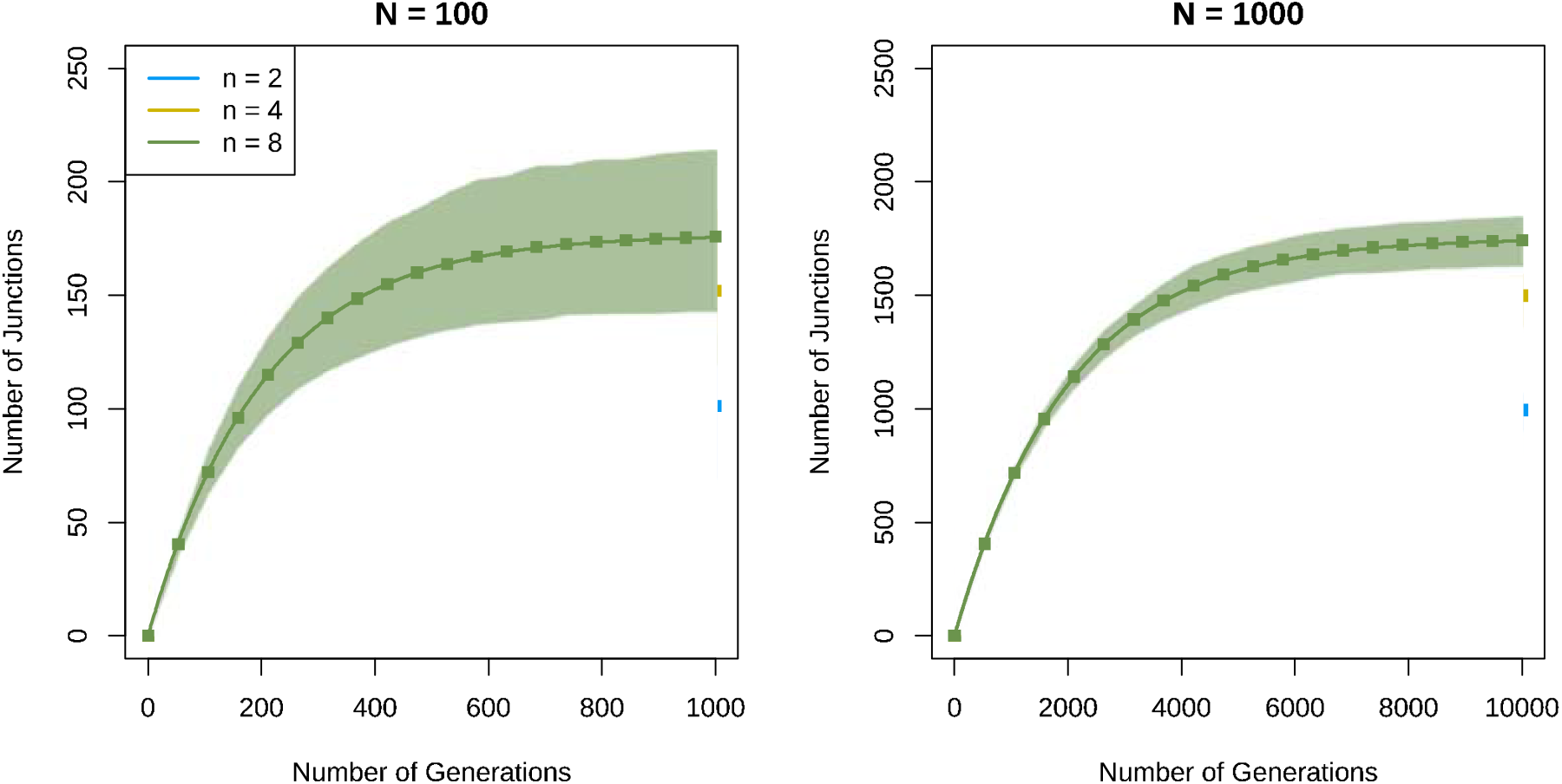
Accumulation of junctions for different numbers of ancestors (2, 4 and 8 unique ancestors), for a small population (N = 100) and a population of intermediate size (N = 1000). Square dots indicate the mean number of junctions observed across 1000 replicate simulations with the GenomeAdmixR package. Shaded areas indicate the 95% Confidence Interval across 1000 replicates. The solid lines indicate the analytical prediction following equation 1. Mean simulation dynamics follow the analytical prediction very closely.

## DISCUSSION

We have presented here a new R package that uses individual-based simulations to study genomic dynamics following admixture. We have shown how both the *ancestry* and *sequence module* can be used for a general approach and more specifically, we have shown how GenomeAdmixR can be used within an experimental evolution framework and how results obtained using GenomeAdmixR are in line with theoretical expectations.

We expect GenomeAdmixR to be applicable in many fields. Firstly, we expect GenomeAdmixR to allow for improved ease of use of individual-based simulations to study patterns in the formation of contiguous tracts of ancestry, and their breakdown due to recombination. The study of the breakdown of ancestry blocks has recently attracted renewed interest, with studies expanding the theory of junctions towards potential marker effects (Janzen *et al*. 2018), studies utilizing patterns in the distribution of ancestry tracts to infer the onset of hybridization (Ungerer *et al*. 1998; Buerkle & Rieseberg 2008; Pool & Nielsen 2009; Gravel 2012; Liang & Nielsen 2014; Corbett-Detig & Nielsen 2017; Medina *et al*. 2018; Schumer *et al*. 2020; Shchur *et al*. 2020) and the inclusion of ancestral tract information in the analysis of human history (Hellenthal *et al*. 2014; Payseur & Rieseberg 2016). GenomeAdmixR complements these analyses by providing a framework to verify findings through simulation. Furthermore, preliminary sequencing data can be used to verify findings using the *sequence module*, providing an additional tool to explore the efficacy of ancestry tract information.

Secondly, we expect GenomeAdmixR to be an important tool in investigating population genomics and designing *E&R* experiments, where GenomeAdmixR can be used to test whether the setup of an experiment is sufficient to detect signatures of selection on a background of existing LD. The functionality of the package allows simulating the effect of *i)* previous population dynamics that occurred before sampling (*natural populations*), *ii)* initial parameters when obtaining the starting population for selection (*mixed populations*) as well as *iii)* variation in genomic and *selection* parameters. Here, we have demonstrated how to use GenomeAdmixR to estimate the level of resolution obtained when combining inbred lines from different populations in the starting population of an E&R experiment (Parts *et al*. 2011; Koide *et al*. 2012). We found that although this model narrowed down the haplotype blocks around the selected maker substantially, this still lacks resolution when compared with traditional sampling (Kofler & Schlötterer 2014; Schlötterer *et al*. 2015) from a single population (even after 30 admixing generations). However, the obtained resolution seems deep enough to investigate questions involving variation between evolutionarily independent entities if these are not possible to address by sampling a single population. For example, Comeault & Matute (2018) used this model to study genomic trajectories following species hybridization.

Thirdly, we expect GenomeAdmixR to function as a useful teaching tool, where it can be used to demonstrate the impact of hybridization on linkage patterns, the subsequent interaction between selection and drift and more generally to provide an easy toolkit to explore the interaction between recombination and hybridization.

GenomeAdmixR shares many similarities with its predecessors SliM (Haller & Messer 2017, 2018) and SELAM (Corbett-Detig & Jones 2016), both powerful admixture simulation programs. However, SLiM is mainly focused on more advanced population demographic scenarios and is not directly suited for tracking ancestry. In contrast, SELAM tracks local ancestry, but requires local compilation of C++ code (which can be difficult) and uses complicated input tables to parameterize the simulations. GenomeAdmixR aims to alleviate both these issues, firstly GenomeAdmixR is specifically focused on tracking ancestry, and its admixture over time. Secondly, because GenomeAdmixR is an R package, parameterization can be done through the use of the R scripting language, which facilitates sharing of code between platforms, replication of approaches and ease of use.

By providing both an *ancestry* and *sequence module*, we think that GenomeAdmixR provides a very complete package that can be used both to explore theoretical expectations (*ancestry module*), and verify these expectations using sequencing data (*sequence module*). Furthermore, because GenomeAdmixR readily accepts sequencing data in popular formats, It can be readily incorporated in existing pipelines. Thus, we expect that GenomeAdmixR has a bright future ahead, with a myriad of potential implementations across different fields of population genetics and molecular ecology.

## AUTHOR CONTRIBUTIONS

TJ and FD conceived the model, TJ developed the R code, FD and TJ jointly wrote and revised the manuscript.

## DATA AVAILABILITY

The R package is available via CRAN on >ttps://cran.r-project.org/package=GenomeAdmixR. All code used for the paper can be found in the Supplemental Material.

## ACKNOWLEDGEMENTS

We thank Omer Markovitch, Pratik R. Gupte and Cyrus Mallon for helpful comments on an earlier version of the manuscript. The authors indicate that they have no conflicts of interest.

## REFERENCES

Abbott, R., Albach, D., Ansell, S., Arntzen, J.W., Baird, S.J.E., Bierne, N., Boughman, J., Brelsford, a., Buerkle, C. a., Buggs, R., Butlin, R.K., Dieckmann, U., Eroukhmanoff, F., Grill, a., Cahan, S.H., Hermansen, J.S., Hewitt, G., Hudson, a. G., Jiggins, C., Jones, J., Keller, B., Marczewski, T., Mallet, J., Martinez-Rodriguez, P., Möst, M., Mullen, S., Nichols, R., Nolte, a. W., Parisod, C., Pfennig, K., Rice, a. M., Ritchie, M.G., Seifert, B., Smadja, C.M., Stelkens, R., Szymura, J.M., Väinölä, R., Wolf, J.B.W. & Zinner, D. (2013). Hybridization and speciation. Journal of Evolutionary Biology, 26, 229–246.

Arnold, M.L. & Kunte, K. (2017). Adaptive Genetic Exchange: A Tangled History of Admixture and Evolutionary Innovation. Trends in Ecology & Evolution, 32, 601–611.

Barghi, N. & Schlötterer, C. (2019). Shifting the paradigm in evolve and resequence studies: From analysis of single nucleotide polymorphisms to selected haplotype blocks. Molecular Ecology, 28, 521–524.

Buerkle, C.A. & Rieseberg, L.H. (2008). The rate of genome stabilization in homoploid hybrid species. Evolution, 62, 266–275.

Burke, M.K. & Rose, M.R. (2009). Experimental evolution with Drosophila. Am J Physiol Regul Integr Comp Physiol.

Burny, C., Nolte, V., Nouhaud, P., Dolezal, M., Schlötterer, C. & Baer, C. (2020). Secondary Evolve and Resequencing: An Experimental Confirmation of Putative Selection Targets without Phenotyping. Genome Biology and Evolution, 12, 151–159.

Chafin, T.K. & Douglas, M.R. (2020). Genome-wide local ancestries discriminate homoploid hybrid speciation from secondary introgression in the red wolf (CanidaelZ: Canis rufus). 1–49.

Comeault, A.A. & Matute, D.R. (2018). Genetic divergence and the number of hybridizing species affect the path to homoploid hybrid speciation. Proceedings of the National Academy of Sciences of the United States of America, 115, 9761–9766.

Corbett-Detig, R. & Jones, M. (2016). SELAM: Simulation of epistasis and local adaptation during admixture with mate choice. Bioinformatics, 32, 3035– 3037.

Corbett-Detig, R. & Nielsen, R. (2017). A Hidden Markov Model Approach for Simultaneously Estimating Local Ancestry and Admixture Time Using Next Generation Sequence Data in Samples of Arbitrary Ploidy. PLoS Genetics, 13, 1–40.

Cottin, A., Penaud, B., Glaszmann, J.C., Yahiaoui, N. & Gautier, M. (2020). Simulation-based evaluation of three methods for local ancestry deconvolution of non-model crop species genomes. G3: Genes, Genomes, Genetics, 10, 569–579.

Crow, J.F. & Kimura, M. (1970). An Introduction to Population Genetics Theory. Harper & Row, New York.

Dennenmoser, S., Schatz, M.C., Sedlazeck, F.J., Zytnicki, M., Nolte, A.W. & Altmüller, J. (2019). Genome-wide patterns of transposon proliferation in an evolutionary young hybrid fish. Molecular Ecology, 0–2.

Eddelbuettel, D. & Francois, R. (2011). Seamless R and C++ integration with Rcpp. Journal of Statistical Software, 40, 1–18. Retrieved from http://www.jstatsoft.org/v40/i08/

Fisher, R.A. (1954). A fuller theory of “Junctions” in inbreeding. Heredity, 8, 187– 197.

Fisher, R.A. (1959). An algebraically exact examination of junction formation and transmission in parent-offspring inbreeding. Heredity, 13, 179–186.

Franssen, S.U., Barton, N.H. & Schlotterer, C. (2017). Reconstruction of haplotype-blocks selected during experimental evolution. Molecular Biology and Evolution, 34, 174–184.

Gerrish, P.J., Colato, A., Perelson, A.S. & Sniegowski, P.D. (2007). Complete genetic linkage can subvert natural selection. PNAS, 104, 6266–6271.

Goudet, J. (2005). HIERFSTAT, a package for R to compute and test hierarchical F -statistics. Molecular Ecology Notes, 2, 184–186.

Gravel, S. (2012). Population genetics models of local ancestry. Genetics, 191, 607–619.

Haller, B.C. & Messer, P.W. (2017). SLiM 2: Flexible, interactive forward genetic simulations. Molecular Biology and Evolution, 34, 230–240.

Haller, B.C. & Messer, P.W. (2018). SLiM 3: Forward Genetic Simulations Beyond the Wright–Fisher Model. Molecular Biology and Evolution, 36, 632–637.

Hellenthal, G., Busby, G.B.J., Band, G., Wilson, J.F., Capelli, C., Falush, D. & Myers, S. (2014). A Genetic Atlas of Human. Science, 343, 747–751.

Huang, W., Massouras, A., Inoue, Y., Peiffer, J., Ràmia, M., Tarone, A.M., Turlapati, L., Zichner, T., Zhu, D., Lyman, R.F., Magwire, M.M., Blankenburg, K., Carbone, M.A., Chang, K., Ellis, L.L., Fernandez, S., Han, Y., Highnam, G., Hjelmen, C.E., Jack, J.R., Javaid, M., Jayaseelan, J., Kalra, D., Lee, S., Lewis, L., Munidasa, M., Ongeri, F., Patel, S., Perales, L., Perez, A., Pu, L.L., Rollmann, S.M., Ruth, R., Saada, N., Warner, C., Williams, A., Wu, Y.Q., Yamamoto, A., Zhang, Y., Zhu, Y., Anholt, R.R.H., Korbel, J.O., Mittelman, D., Muzny, D.M., Gibbs, R.A., Barbadilla, A., Johnston, J.S., Stone, E.A., Richards, S., Deplancke, B. & Mackay, T.F.C. (2014). Natural variation in genome architecture among 205 Drosophila melanogaster Genetic Reference Panel lines. Genome Research, 24, 1193–1208.

Janzen, T., Nolte, A.W. & Traulsen, A. (2018). The breakdown of genomic ancestry blocks in hybrid lineages given a finite number of recombination sites. Evolution, 72, 735–750. Retrieved from http://doi.wiley.com/10.1111/evo.13436

Kawecki, T.J., Lenski, R.E., Ebert, D., Hollis, B., Olivieri, I. & Whitlock, M.C. (2012). Experimental evolution. Trends in ecology & evolution, 27, 547–60.

Keeling, P.J. & Palmer, J.D. (2008). Horizontal gene transfer in eukaryotic evolution. Nature Reviews Genetics, 9, 605–618.

Kessner, D. & Novembre, J. (2014). Forqs: Forward-in-time simulation of recombination, quantitative traits and selection. Bioinformatics, 30, 576– 577.

Kessner, D. & Novembre, J. (2015). Power analysis of artificial selection experiments using efficient whole genome simulation of quantitative traits. Genetics, 199, 991–1005.

Kofler, R. & Schlötterer, C. (2014). A guide for the design of evolve and resequencing studies. Molecular Biology and Evolution, 31, 474–483.

Koide, T., Goto, T. & Takano-Shimizu, T. (2012). Genomic mixing to elucidate the genetic system of complex traits. Experimental Animals, 61, 503–509.

Lavretsky, P., Janzen, T. & McCracken, K.G. (2019). Identifying hybrids & the genomics of hybridization: Mallards & American black ducks of Eastern North America. Ecology and Evolution, 9, 3470–3490.

Leitwein, M., Gagnaire, P.A., Desmarais, E., Berrebi, P. & Guinand, B. (2018). Genomic consequences of a recent three-way admixture in supplemented wild brown trout populations revealed by local ancestry tracts. Molecular Ecology, 27, 3466–3483.

Liang, M. & Nielsen, R. (2014). The lengths of admixture tracts. Genetics, 197, 953–967.

MacKay, T.F.C., Richards, S., Stone, E.A., Barbadilla, A., Ayroles, J.F., Zhu, D., Casillas, S., Han, Y., Magwire, M.M., Cridland, J.M., Richardson, M.F., Anholt, R.R.H., Barrón, M., Bess, C., Blankenburg, K.P., Carbone, M.A., Castellano, D., Chaboub, L., Duncan, L., Harris, Z., Javaid, M., Jayaseelan, J.C., Jhangiani, S.N., Jordan, K.W., Lara, F., Lawrence, F., Lee, S.L., Librado, P., Linheiro, R.S., Lyman, R.F., MacKey, A.J., Munidasa, M., Muzny, D.M., Nazareth, L., Newsham, I., Perales, L., Pu, L.L., Qu, C., Ràmia, M., Reid, J.G., Rollmann, S.M., Rozas, J., Saada, N., Turlapati, L., Worley, K.C., Wu, Y.Q., Yamamoto, A., Zhu, Y., Bergman, C.M., Thornton, K.R., Mittelman, D. & Gibbs, R.A. (2012). The Drosophila melanogaster Genetic Reference Panel. Nature, 482, 173–178.

Macleod, A.K., Haley, C.S., Woolliams, J.A. & Stam, P. (2005). Marker densities and the mapping of ancestral junctions. Genetical Research, 85, 69–79.

Medina, P., Thornlow, B., Nielsen, R. & Corbett-Detig, R. (2018). Estimating the timing of multiple admixture pulses during local ancestry inference. Genetics, 210, 1089–1107.

Orozco-Terwengel, P., Kapun, M., Nolte, V., Kofler, R., Flatt, T. & Schlãtterer, C. (2012). Adaptation of Drosophila to a novel laboratory environment reveals temporally heterogeneous trajectories of selected alleles. Molecular Ecology, 21, 4931–4941.

Otte, K.A. & Schlötterer, C. (2020). Detecting selected haplotype blocks in Evolve and Resequence experiments. Molecular Ecology Resources, 1–37.

Parts, L., Cubillos, F.A., Warringer, J., Jain, K., Salinas, F., Bumpstead, S.J., Molin, M., Zia, A., Simpson, J.T., Quail, M.A., Moses, A., Louis, E.J., Durbin, R. & Liti, G. (2011). Revealing the genetic structure of a trait by sequencing a population under selection. 1131–1138.

Payseur, B.A. & Rieseberg, L.H. (2016). A genomic perspective on hybridization and speciation. Molecular Ecology, 25, 2337–2360.

Pool, J.E. & Nielsen, R. (2009). Inference of historical changes in migration rate from the lengths of migrant tracts. Genetics, 181, 711–719.

Schlötterer, C., Kofler, R., Versace, E., Tobler, R. & Franssen, S.U. (2015). Combining experimental evolution with next-generation sequencing: A powerful tool to study adaptation from standing genetic variation. Heredity, 114, 431–440.

Schumer, M., Cui, R., Powell, D.L., Dresner, R., Rosenthal, G.G. & Andolfatto, P. (2014a). High-resolution mapping reveals hundreds of genetic incompatibilities in hybridizing fish species. eLife, 2014, 1–21.

Schumer, M., Powell, D.L. & Corbett-Detig, R. (2020). Versatile simulations of admixture and accurate local ancestry inference with mixnmatch and ancestryinfer. Molecular Ecology Resources, 20, 1141–1151.

Schumer, M., Rosenthal, G.G. & Andolfatto, P. (2014b). How common is homoploid hybrid speciation? Evolution, 68, 1553–1560.

Schumer, M., Xu, C., Powell, D.L., Durvasula, A., Skov, L., Holland, C., Blazier, C., Sankararaman, S., Andolfatto, P., Rosenthal, G.G. & Przeworski, M. (2018). Natural selection interacts with recombination to shape the evolution of hybrid genomes. Science, 360, 656–660.

Shchur, V., Svedberg, J., Medina, P., Corbett-Detig, R. & Nielsen, R.A.S.M. (2020). On the distribution of tract lengths during adaptive introgression. G3: Genes, Genomes, Genetics, 10, 3663–3673.

Stam, P. (1980). The distribution of the fraction of the genome identical by descent in finite random mating populations. Genetical Research, 35, 131.

Team, R.C. (2020). R: A Language and Environment for Statistical Computing.

Tobler, R., Franssen, S.U., Kofler, R., Orozco-terwengel, P., Nolte, V., Hermisson, J. & Schlotterer, C. (2013). Massive Habitat-Specific Genomic Response in D. melanogaster Populations during Experimental Evolution in Hot and Cold Environments. Molecular Biology and Evolution.

Turner, T.L. & Miller, P.M. (2012). Investigating natural variation in drosophila courtship song by the evolve and resequence approach. Genetics, 191, 633– 642.

Ungerer, M.C., Baird, S.J., Pan, J. & Rieseberg, L.H. (1998). Rapid hybrid speciation in wild sunflowers. Proceedings of the National Academy of Sciences of the United States of America, 95, 11757–11762.

Vlachos, C. & Kofler, R. (2018). MimicrEE2: Genome-wide forward simulations of Evolve and Resequencing studies. PLoS Computational Biology, 14, 1–10. Retrieved from http://dx.doi.org/10.1371/journal.pcbi.1006413

Weir, B.S. & Cockerham, C.C. (1984). Estimating F-Statistics for the Analysis of Population Structure Author (s): B. S. Weir and C. Clark Cockerham Published bylZ: Society for the Study of Evolution Stable URLlZ: http://www.jstor.org/stable/2408641. Evolution, 38, 1358–1370.

Wickham, H. (2009). ggplot2: Elegant Graphics for Data Analysis - Hadley Wickham - Google Books. Retrieved September 9, 2020, from https://books.google.nl/books?hl=nl&lr=&id=XgFkDAAAQBAJ&oi=fnd&pg=PR8&dq=ggplot2&ots=so58bP5WbN&sig=ThS6gEgxaK9XADL_HeG5gDU04Pc#v=onepage&q=ggplot2&f=false

Zhou, Y., Qiu, H. & Xu, S. (2017). Modeling Continuous Admixture Using Admixture-Induced Linkage Disequilibrium. Scientific Reports, 7, 1–10.

