## Supplement for "Individual-based simulations of genome evolution with ancestry: the GenomeAdmixR R package"

*Evolve and resequence - E&R experiments*

One of the goals in *E&R* experiments is to identify underlying genetic loci driving responses to selection (Burke & Rose 2009; Schlötterer *et al.* 2015). Candidate loci are investigated by comparing genomic changes in samples sequenced before and resequenced after selection (Long *et al.* 2015). Considering that experimental selection can take a long time in the lab, the ability to simulate genomic dynamics under different scenarios of selection becomes pivotal to take informed decisions. Among one of the several decisions that researchers must take to improve experimental designs is how to maximize the initial genetic variation. Increased genetic variability in the starting population improves the chances of capturing additional candidate loci, but Linkage Disequilibrium (LD) can mask the changes at individual positions (Kessner & Novembre 2015). GenomeAdmixR can help make decisions by comparing alternative sampling scenarios for E&R experiments, using individual-based simulations.

*The comparison: alternative sampling scenarios for E&R*

The most common sampling method used in *E&R* experiments starts with a “w*ell* *mixed population*” founded by individuals sampled from a single “*natural population”* (***Model 1***). This method minimizes LD while trying to compensate for genetic variability by sampling a large number of individuals (Kofler & Schlötterer 2014)*.* Alternatively, genetic variability can be maximized by sampling from different populations (***Model 2***), but this generates additional *LD* (Kawecki *et al.* 2012). Massive hitchhiking genomic regions that arise with *LD* can prevent the ability to differentiate target loci from the genome background (Kofler & Schlötterer 2014; Schlötterer *et al.* 2015). Given that inbred lines with interesting phenotypes are often available for model species, it has been proposed that genetic bases of populations or species differences can be investigated using *E&R* (Parts *et al.* 2011) as long as the starting population is mixed well enough (as for *QTL* studies). This method has been used in yeasts to fine tune *QTL* studies from phenotypically extreme inbred lines using experimental selection (Parts *et al.* 2011; Koide, Goto & Takano-Shimizu 2012). Whether the *LD* is broken down enough (*Model 2*) compared with traditional sampling in *E&R* (*Model 1*), or what level of genomic resolution is expected if implemented in animal species are questions that we can address by exploring their genomic responses to simulated selection. Here we performed simulations in three blocks (the workflow is summarized in Figure 2): 1) Simulation of two ‘*natural populations*” from which to sample individuals; then , 2) Simulation of two alternative “*mixed populations*” in the lab (*Models 1 and 2*); And, 3) Simulation of an *E&R* experiment with selection. Selection is performed on a single locus using the same selection regimen. We compared the performance of assessed models and ultimately their capacity to detect truly selected loci from the genome background.

1) *Using GenomeAdmixR to simulate natural populations*

The two *natural populations* (*Population 1* and *Population 2*) were simulated by an allopatric split of an initial *source population* (N=3000) (function simulate_admixture). *Natural populations* continue to diverge until they reach high genetic differentiation (average F_ST_ ~ 0.4) across the genome (function simulate_admixture_until). High genetic differentiation between the two populations is simulated to test the effect of combining highly divergent populations to perform experimental evolution.

2) *Using GenomeAdmixR to simulate mixed populations in the lab*

Instead of individuals, we simulated the sampling of isofemale lines [a common sampling method in *Drosophila* and other insects [*e.g.* (David *et al.* 2005; Burny *et al.* 2020) to demonstrate additional utilities of the package (function create_isofemale_line). In ***Model 1*,** the *mixed population* is created by admixing 100 isofemale lines sampled from *Population 1*. This population reproduces in the lab for two generations before the selection experiment, because the model takes advantage of the minimal *LD* when sampling a single population. In ***Model 2***, the *mixed population* is generated by only two isofemales sampled from two different *natural populations*. The new population is kept in the lab for an additional 30 generations before the selection experiment, because LD is massive in the first generations. This approach is in line with Quantitative Trait Loci (QTL) studies.

3) *Using GenomeAdmixR to simulate an E&R with selection*

To implement selection (function *simulate_admixture*), we need to define *i)* the selection matrix (markers and selection coeficients), *ii)* initial allele frequencies (*IAF*), and *iii)* the total number of tracking markers. These parameters were defined in a 0.2 Morgan genome window, with the rest of chromosome evolving neutrally. The window is used to track not only the results of selection on a single locus, but also hitchhiking effects on surrounding areas (300 markers). We used the same selection matrix on a single marker for each model (w ~ aa=0.5, Aa=0.5, AA=1.0), and simulated selection for 20 generations, a time period commonly used in these experiments (Orozco-Terwengel *et al.* 2012). Because the response to selection is highly influenced by the initial gene pool in the starting *mixed population*, the target markers were chosen based on similar initial allele frequencies (IAF) [0.1 > *IAF <*0.2 using the function calculate_marker_frequency]. The genomic positions of selected makers are not exactly the same for each model, but the mentioned conditions as well as the genomic window are the same (0.2 Morgan).

*Results*

We investigated genome-wide responses to the simulated *E&R* experiment with selection by estimating allele frequency changes (Δ allele fq = fq after – fq before selection) after 20 generations of selection (300 tracked markers). Both models showed a clear peak in allele frequency that increase over time around the truly selected marker (Figure 3). The relative differences in *LD* can be appreciated in the size of the responding regions (hitchhiking regions). The resolution to detect the truly selected marker was much higher in *Model 1* vs *Model 2* (Figure 3). The Δ allele frequency at the truly selected marker was much easier to distinguish from background variation in *Model 1*. A ~ 0.08 Morgan region seems to respond to selection as a result of increased hitchhiking areas in *Model 2*. The alternative model to investigate variation between population or species (*Model 2*) does not seem to provide as good a resolution as traditional sampling (*Model 1*). However, the obtained results show a relatively narrowed haplotype structure compared with initial massive *LD* when combining different entities. Model 2 can potentially work as long as the question of interest involves lines that have evolved independently (*e.g.* extreme phenotypes between populations), although with lower resolution that will require additional tuning.

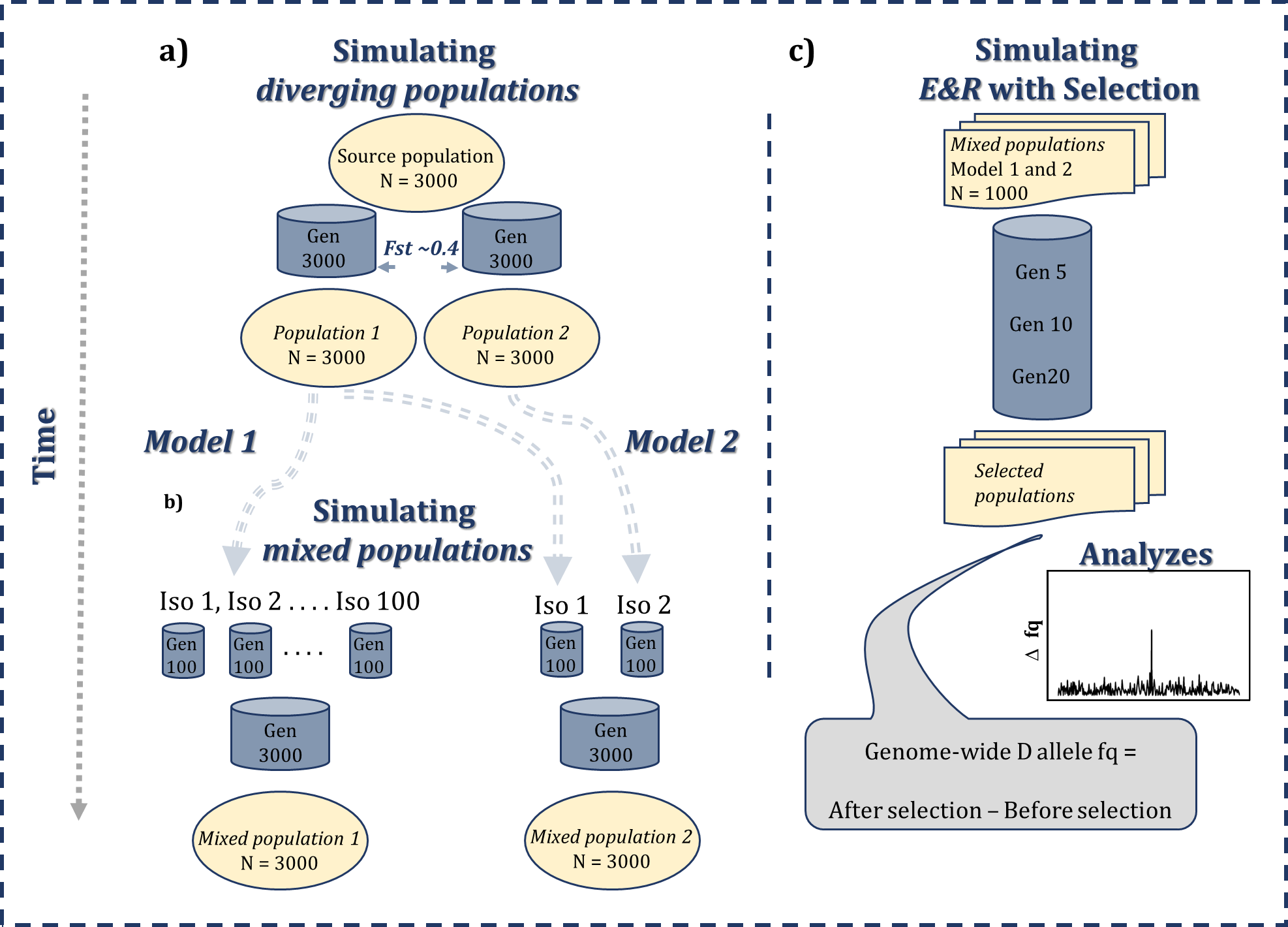
**Figure S1. Workflow used to compare two sampling models to create *mixed populations* for *E&R* experiments using with the GenomeAdmixR package. a)** The simulation starts with an original source population that diverge into two populations. Then, **b)** Isofemales lines are sampled to created mixed populations either with 100 lines from a single population (***Model 1***) or two lines from different populations (***Model 2*)**. **c)** Genome-wide responses to selection on a single marker are then compared between mixed populations of the two models (starting allele frequency ~ 0.1). The effect of selection is analyzed by tracking changes of allele frequency in 300 biallelic markers around the selected marker.

*Match with theory*

The GenomeAdmixR package allows for the simulation of the breakdown of contiguous blocks of ancestry due to recombination. From the extended theory of junctions (Janzen, Nolte & Traulsen 2018) we know that the expected number of junctions (where a junction delineates the end of one contiguous stretch of genomic content from the same ancestor, and the start of another), depends on the initial heterozygosity $H_{0}$ (where heterozygosity here reflects a locus with two alleles stemming from two different ancestors, e.g. hetero-ancestry, which is mathematically identical assuming no mutation), such that:

$E\left[ J_{t} \right]={2NH}_{0}-2NH_{0}\left( 1-\frac{1}{2N} \right)^{t}$ (1)

where *N* is the population size. Janzen’s extended theory of junctions focuses specifically on the scenario with two ancestors, where *H_0_* *= 2pq*, where *p* is the frequency of one ancestor, and *q = 1-p* the frequency of the other ancestor in the ancestral swarm. GenomeAdmixR allows for an arbitrary number of ancestors. If we for simplicity assume an even distribution in the ancestral swarm, such that the frequency of ancestor *i* is $\frac{1}{i}$, then, the initial heterozygosity is given by $H_{0}=1-\sum_{i=1}^{n} \frac{1}{i^{2}}=1-\frac{1}{n}$, where *n* indicates the number of ancestors at *t = 0*. If equation (1) can be generalized to the multiple ancestor case, we expect the accumulation of junctions in GenomeAdmixR simulations to follow equation (1). We simulate the accumulation of junctions using GenomeAdmixR for two population sizes (100 and 1000 individuals) and vary the number of ancestors *n* in [2, 4, 8]. We find that when we average the number of accumulated junctions over 1000 replicates, the average number of junctions closely follows that of our analytical expectation (Figure S3 and Figure 4 in the main text).

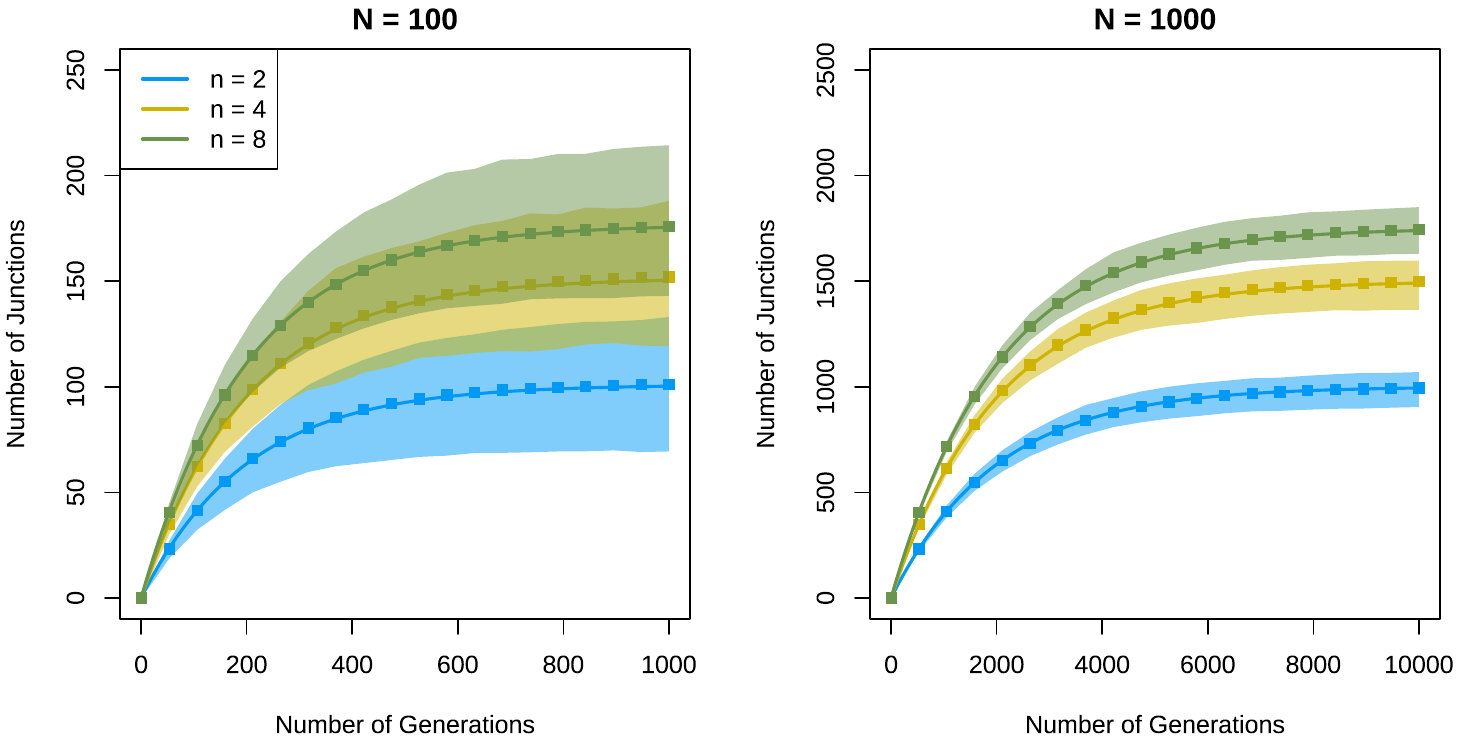

Figure S3 / 4. Identical to Figure 4 in the main text, added for quick reference. Accumulation of junctions for different numbers of ancestors (2, 4 and 8 unique ancestors), for a small population (N = 100) and a population of intermediate size (N = 1000). Square dots indicate the mean number of junctions observed across 1000 replicate simulations with the GenomeAdmixR package. Shaded areas indicate the 95% Confidence Interval across 1000 replicates. The solid lines indicate the analytical prediction following equation 1. Mean simulation dynamics follow the analytical prediction very closely.

*Average heterozygosity*

The expected heterozygosity after *t* generations, is given by: $H_{0}\left( 1-\frac{1}{2N} \right)^{t}$, where $H_{0}$ is the initial heterozygosity at *t = 0*. Assuming even proportions of ancestors, $H_{0}=1-\frac{1}{n}$, and we can formulate an expectation of the average heterozygosity, given any number of founders. Simulations closely match these results. Here, we tested whether the average heterozygosity follows our expectations for two scenarios: either with 2 ancestors and a population size of 1000 individuals, or with 4 ancestors and a population size of 100 individuals. Both simulations were run for 1500 generations, and every 25 generations heterozygosity was measured. We performed 10 replicates per scenario. Across both scenarios, we find that the median result across the replicates closely follows the theoretical expectation (Figure S5).

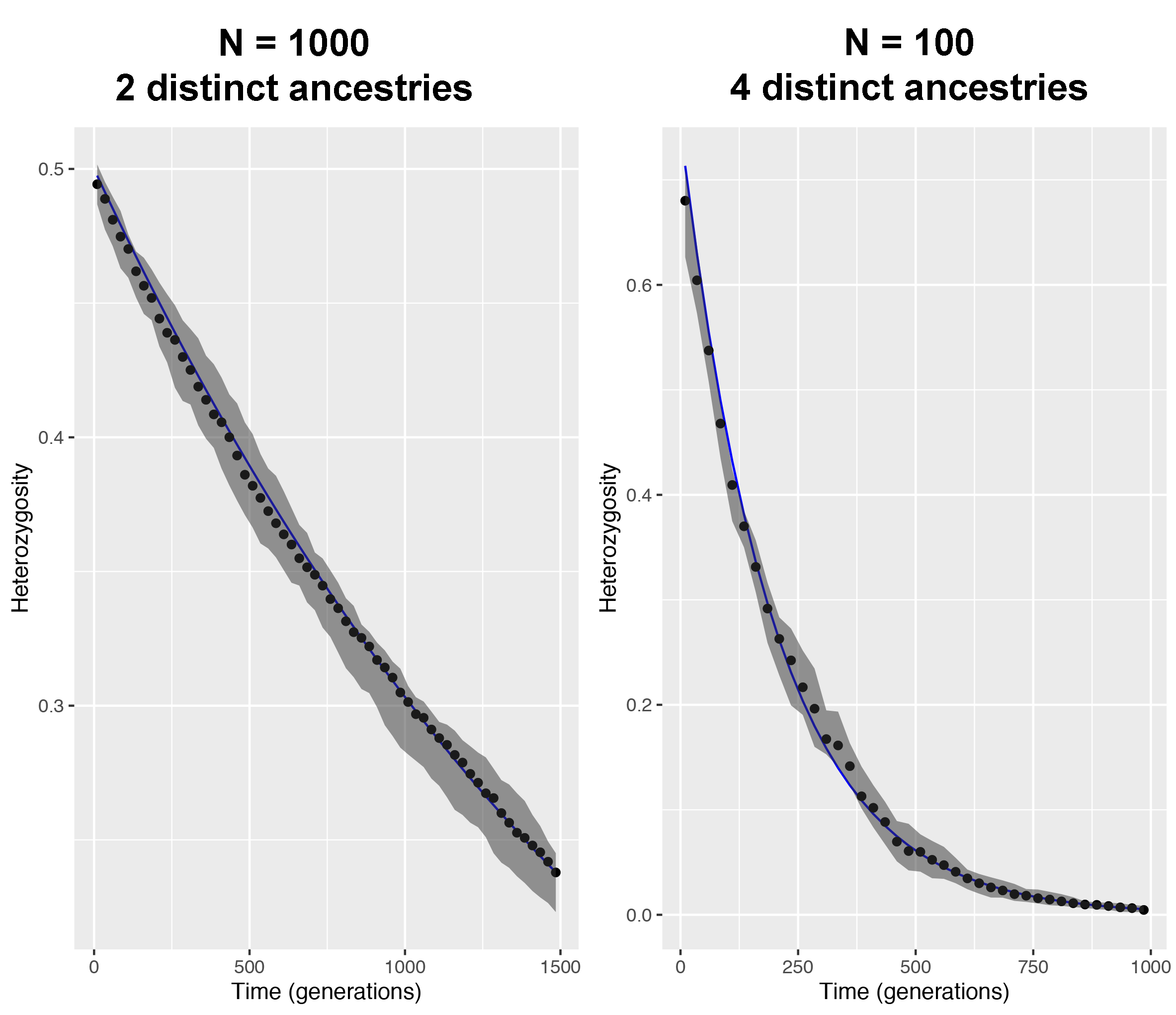

**Figure S5.** Average heterozygosity for simulations (dots indicate the median across 10 replicates, grey area indicating 95% CI) run with either a population size of 1000 individuals and 4 ancestors (left panel) or 100 individuals and 2 ancestors (right panel). The blue line indicates the expected heterozygosity.

REFERENCES

Burke, M.K. & Rose, M.R. (2009) Experimental evolution with Drosophila. *Am J Physiol Regul Integr Comp Physiol*.

Burny, C., Nolte, V., Nouhaud, P., Dolezal, M., Schlötterer, C. & Baer, C. (2020) Secondary Evolve and Resequencing: An Experimental Confirmation of Putative Selection Targets without Phenotyping. *Genome Biology and Evolution*, **12**, 151–159.

David, J.R., Gibert, P., Legout, H., Pétavy, G., Capy, P. & Moreteau, B. (2005) Isofemale lines in Drosophila: an empirical approach to quantitative trait analysis in natural populations. *Heredity*, **94**, 3–12.

Janzen, T., Nolte, A.W. & Traulsen, A. (2018) The breakdown of genomic ancestry blocks in hybrid lineages given a finite number of recombination sites. *Evolution*, **72**, 735–750.

Kawecki, T.J., Lenski, R.E., Ebert, D., Hollis, B., Olivieri, I. & Whitlock, M.C. (2012) Experimental evolution. *Trends in ecology & evolution*, **27**, 547–60.

Kessner, D. & Novembre, J. (2015) Power analysis of artificial selection experiments using efficient whole genome simulation of quantitative traits. *Genetics*, **199**, 991–1005.

Kofler, R. & Schlötterer, C. (2014) A guide for the design of evolve and resequencing studies. *Molecular Biology and Evolution*, **31**, 474–483.

Koide, T., Goto, T. & Takano-Shimizu, T. (2012) Genomic mixing to elucidate the genetic system of complex traits. *Experimental Animals*, **61**, 503–509.

Long, A., Liti, G., Luptak, A. & Tenaillon, O. (2015) Elucidating the molecular architecture of adaptation via evolve and resequence experiments. *Nature Reviews Genetics*, **16**, 567–582.

Orozco-Terwengel, P., Kapun, M., Nolte, V., Kofler, R., Flatt, T. & Schlãtterer, C. (2012) Adaptation of Drosophila to a novel laboratory environment reveals temporally heterogeneous trajectories of selected alleles. *Molecular Ecology*, **21**, 4931–4941.

Parts, L., Cubillos, F.A., Warringer, J., Jain, K., Salinas, F., Bumpstead, S.J., Molin, M., Zia, A., Simpson, J.T., Quail, M.A., Moses, A., Louis, E.J., Durbin, R. & Liti, G. (2011) Revealing the genetic structure of a trait by sequencing a population under selection. , 1131–1138.

Schlötterer, C., Kofler, R., Versace, E., Tobler, R. & Franssen, S.U. (2015) Combining experimental evolution with next-generation sequencing: A powerful tool to study adaptation from standing genetic variation. *Heredity*, **114**, 431–440.
